# Prevalence Of Multi Drug Resistant Bacteria On Environmental And Medical Device Surfaces Of Korea Nepal Friendship Hospital

**DOI:** 10.1101/2022.02.20.481232

**Authors:** Nisha Paudel, Nagendra Prasad Awasthi, Binod Lekhak, Amrit Acharya

**Affiliations:** CCMB; Deakin University; Tribhuvan University; Padma Kanya Multiple Campus

**Keywords:** Disk diffusion, Environmental monitoring, MDR, Nosocomial infections, assay

## Abstract

The microbial monitoring of environmental and medical-devices’ surface is used to evaluate efficacy of routine cleaning and disinfection practises and to detect the presence of specific Nosocomial Pathogens. The prevalence of Multidrug Resistance organisms in hospital premises projects serious problems in transmitting to susceptible hosts which is difficult to treat. A cross sectional descriptive research was conducted from December 2016 to June 2017 at the pathology laboratory of Korea Nepal Friendship Hospital (KNFH). A total 140 samples were considered, encompassing the medical devices of the hospital (100), housekeeping surfaces (15) and air (25). Susceptibility test for bacterial isolates was done by disk diffusion assay. Of the total 140 samples taken and analysed, 100% showed growth positivity. In most of the swabs taken, Coagulase Negative Staphylococci was dominant, followed by *Staphylococcus aureus, Streptococcus* spp. *Micrococcus* spp., *E coli, Pseudomonas* spp., *Bacillus* spp., *Acinetobacter* spp., *Klebsiella* spp., Fungi, and least *Proteus* spp. The dry surfaces were dominantly contaminated by gram positive bacteria whereas moistened surfaces like wash basin were contaminated by gram negative as well as gram positive bacteria. Total 277 strains were exposed to various class of antibiotics, among the gram positive environmental isolates, Coagulase Negative Staphylococci 16 (34.78%) had highest MDR prevalence followed by *Staphylococcus aureus* 8 (29.62%), *Streptococcus* spp. 4 (12.90%), *Micrococcus* spp. 4 (9.30%) and no MDR was shown by any *Bacillus* spp isolates. Whereas, in case of gram negative, *Klebsiella* spp. 6 (35.29%) had highest MDR prevalence

followed by *Acinetobacter* spp. 6 (31.57%), *E. coli* 8 (27.58%), *Pseudomonas* spp. 4 (18.18%), and lastly *Proteus* spp. with no MDR at all. The thick dirt covering the cotton swabs and heavy microbial load on them has displayed not only disinfecting practice but also cleaning practice is missing. Heavy contamination shows possible NIs breakout, it’s important to have routine microbial assessment with standard protocol and find ways to decrease its load.

## 1. INTRODUCTION AND OBJECTIVES

### 1.1 Background

Environmental surfaces in Healthcare centers act as reservoirs for bacteria and can as well serve as vectors of the bacterial pathogens (Boyce et al 1997 and Otter et al 2013). Environmental surfaces include the floor surface, door knob, AC and wash basin and medical-devices include the devices like nebulizers, thermometer, Pulse Oximeter (POM) which is directly in contact with the body of the patients. Hands are the main sources, followed by medical devices, such as: catheters, ventilators, endoscopes, sphygmomanometers, autoscopes, thermometers, stethoscopes, computer keyboards etc (Gabriele et al 2014). Although hand hygiene is important to minimize the impact of NIs, cleaning and disinfecting environmental surfaces is fundamental in reducing their potential contribution to the incidence of NIs (CDC 2003).

Environmental surfaces carry the risk of disease transmission but can be safely decontaminated using less rigorous methods than those used on medical instruments and devices (CDC 2003). These areas have high potential of being contaminated by the pathogenic microorganisms as they are in contact of patients and health care workers(HCWs). The surface, therefore, is considered as one of the potential reservoirs for the pathogen, not the only source of exposure. Due to the intensive and /or improper use of antibiotics, the emergence of antibiotic resistant micro-organisms is increasing rapidly around the globe creating a serious threat; many of the pathogens that cause nosocomial infection have a high level of resistance to antibiotic treatment (Pfaller et al 1997). Infections from drug resistant *Escherichia coli, Staphylococcus aureus*, and *Pseudomonas aeruginosa* are becoming common (Oli et al 2013).

Accelerated improvements in diagnostic and therapeutic procedures have helped significant progress in medical era but plentiful using of invasive methods cause many health care problems every day (Oskouie et al 2013). So, hospital has increasingly become unsafe place for patient during their stay. Health care associated infections are important causes of mortality, substantial morbidity, prolonged hospital stays, and increased costs (Fanos et al 2002) and also increases resistance of microorganisms to antimicrobials. According to WHO (2014), every hundred 100 hospitalized patients at any given time, 7 in developed and 10 in developing countries will acquire health care-associated infection. The incidence of NIs is estimated at 5-10% in tertiary care hospitals reaching up to 28% in ICU (Jain et al 2007). This leads into development of new discipline called as SENIC (Study on the Efficacy of Nosocomial Infection Control) which help to assess the surveillance and control activities in hospitals (Hughes 1988).

The development of MDR is usually attributed to the multidrug efflux pump, conjugative R-plasmids, integrons (Stephen et al 2005). Resistance and its spread among bacteria is generally the result of selective antibiotic pressure. (WHO Global Strategy for Containment of Antimicrobial Resistance. 2001). Inappropriate and uncontrolled use of antimicrobial agents including overprescribing, administration of suboptimal doses, insufficient duration of treatment, and misdiagnosis leading to inappropriate choice of drug, contribute to this. In health care settings, the spread of resistant organisms is facilitated when handwashing, barrier precautions, and equipment cleaning are not optimal. The emergence of resistance is also favored by underdoing due to shortage of antibiotics, where lack of microbiological laboratories results in empiric prescribing, and where the lack of alternate agents compounds the risk of therapeutic failure (Panta et al 2011).

According to a literature review of national or multicentre studies published from 1995 to 2008, the overall prevalence of NI in developed countries varies between 5.1% and 11.6% moreover, studies conducted in health-care settings in developing countries report, rates markedly higher than those in developed countries published by WHO in the aricticle “The Burden of Health Care-Associated Infection Worldwide” 2014.

In context of resource poor country like Nepal, it has been found that, about 10.5 % of patients admitted to acute care hospitals develop NIs (Tuladhar et al 1990). In tertiary care hospital of Nepal, the overall point prevalence of NI is reported to be 2.4% (Lamichhane et al 2001).

Korea-Nepal Friendship Hospital (KNFH) was established in 1999. Despite of its long history of establishment and services of this hospital, no microbial investigation for surveillance of NIs and status of MDR has been conducted. So, this study may be an evidence for the need of management of NI and control development of MDR strain. Though, the study was carried out in a single hospital, this study will arrest the attention of all the hospitals’ management committee in making appropriate surveys and investigations to control infections caused by hospital strains.

This study was conducted to assess the status of microbiological flora from the environmental and Medical-Device Surfaces of Korea Nepal Friendship Hospital. The result of this study will help to make plans and policies to prevent NIs and bring awareness about cleaning and disinfection of the environment as well as medical devices in KNFH.

### 1.2 OBJECTIVES

#### 1.2.1 General Objective

To assess the status of Multi Drug Resistant (MDR) bacteria from the Environmental and Medical Device Surfaces of Korea Nepal Friendship Hospital.

#### 1.2.2 Specific Objectives

To enumerate and identify the microbial load present in hospital Environment (air, floor and wash basins)
To enumerate and identify the microbial load present in Medical-Device surfaces of health care center.
To perform antibiotic sensitivity test of bacterial isolates to assess MDR.

## 2. MATERIALS AND METHOD

### 2.1 Materials

All the materials used during study are given in appendix-A.

### 2.2 Methodology

#### 2.2.1 Study design

Cross sectional descriptive hospital based study was conducted to evaluate hospital environment and surfaces of medical devices.

#### 2.2.2 Collaborating institutions

i. Korea Nepal Friendship Hospital
ii. Padma Kanya Multiple Campus

#### 2.2.3 Sampling frame

The environmental samples were air, floor wash basin and medical devices in direct contact with patients in different operation theater, wards and OPDs.

#### 2.2.4 Study period

The present study was conducted from December 2016 to June 2017 at the Pathology laboratory of KNFH, Bhaktapur and microbiology department of Padma Kanya Multiple campus, Kathmandu.

#### 2.2.5 Sample size

A total of 140 samples were collected that includes 100 from swab of medical devices, 15 swabs from housekeeping surfaces and 25 from plate exposure for air sampling.

### 2.3 Laboratory methodology

#### 2.3.1 Collection of sample and processing of the samples

The present work includes the samples such as air, surfaces of medical devices which is in contact of patients and housekeeping surfaces.

##### 2.3.1.1 Air Samples

A total of 25 air samples were collected using gravity settlement method whereby the culture plates, including NA, BA, MA and SDA were exposed to air. The plates were left open with lid by its side at 3-4 feet height from the floor for a time period of 20 minutes. The plates were then incubated at 37°C for 24 hours for bacterial growth and 37°C for 3 to 5 days for fungal growth. Enumeration and observation was done. Isolated bacteria were subcultured into NA and then identified with colony morphology, gram staining and different biochemical tests. Lastly the AST was performed (CDC, 2003)

##### 2.3.1.2 Housekeeping surface samples

A total of 15 surface samples were collected from different wards. The surface samples include samples from floor, AC and wash basin. Surface samples were taken using by sterile cotton swabs dipped in normal saline. For the purpose of quality control each batch of swab was tested for sterility by streaking on culture medium and incubated at 37°C for observing colonies. For proper collection of samples, it was made sure that the swab was taken perpendicularly with only its tip of the swab touching the surface area keeping the area to be swabbed fixed that is 3*3 cm square and streaked into NA, BA and MA. The plates were then incubated at 37°C for 24 hours for bacterial growth. Enumeration and observation was done. Isolates were subcultured into NA and then identified with colony morphology, gram staining and different biochemical tests and AST of the isolates was performed (CDC 2003)

##### 2.3.1.3 Medical devices’ surface swab samples

A total of 104 medical devices’ surface swab samples were collected from different wards and OPDs (Given in appendix B). The surface swab samples include samples from all the medical devices from all operation theaters, wards and OPDs. Surface samples were taken using sterile cotton swabs dipped in normal saline. For the purpose of quality control each batch of swab was tested for sterility by streaking on culture medium and incubated at 37°C for observing colonies. For proper collection of samples, it was made sure that the swab was taken perpendicularly with only tip of the swab touching the surface keeping the area to be swabbed fixed that is 3*3 cm square. and streaked into NA, BA and MA, the plates were then incubated at 37°C for 24 hours for bacterial growth. Enumeration and observation was done. Bacterial isolates were subcultured into NA and then identified with colony morphology, gram staining and different biochemical tests. Finally, the AST was performed (CDC 2003).

#### 2.3.2 Identification of bacterial isolates

Isolated organisms were identified on the basis of colonial characteristics, Gram’s staining and various biochemical tests including Oxidase, Catalase, Sulphide, Indole, Motility, MR, VP, TSI, Citrate, Urease, OF and Coagulase as described by Cheesburgh (2006). Preparation of media, procedure for grams staining and biochemical tests are given in Appendix B, C and D respectively.

#### 2.3.3 Antibiotic susceptibility test (AST)

Antibiotic susceptibility test of all isolates was performed by modified Kirby Bauer disc diffusion method. Procedure for AST is given as:

1. Preparation of medium: Medium used for running AST was Muller Hinton agar. Medium was prepared as per directions provided by manufacturer, care was taken to maintain height of medium while pouring (4mm in 90mm diameter plates i.e. 25ml per plate).
2. Preparation of inoculum: Inoculum was prepared by adding pure colonies of organisms to 5ml Nutrient broth and incubated at 37°C for 4 hours. The prepared inoculum was compared with McFarland tube number 0.5 (Preparation of McFarland tube number 0.5 is given in appendix E.
3. Performing sensitivity test: After proper turbidity was achieved, a new sterile cotton swab was submerged in the suspension, lifted out of the broth, and the excess fluid was removed by pressing and rotating the swab against the wall of the tube. The swab was then used to inoculate the entire surface of the Mueller Hinton agar plate three times, rotating the plate 60 degrees between each inoculation. The inoculum was allowed to dry (usually taking only a few minutes but no longer than 15 minutes) before the discs are placed on the plates. The discs were then placed on the agar with sterile forceps and tapped gently to ensure the adherence to the agar. The plates containing the discs were incubated at 37°C for 24 hours.
4. Observation and result interpretation After overnight incubation, the diameter of each zone of inhibition was measured with a ruler. The ruler was positioned across the center of the disc to make these measurements. The results were recorded in millimeters (mm) and interpretation of susceptibility was obtained by comparing the results to the standard zone interpretative chart provided by the company (APPENDIX F)

### 2.4 Quality control

Quality control was applied in various areas during the study period for the accurate interpretation of results. During the preparation, sterilization, storage and use of media, instructions provided by the manufacturer were strictly followed to avoid alteration of nutritional, selective, inhibitory and biochemical properties of the media. The performance of newly prepared media was tested using the control species of bacteria (i.e., known organisms giving positive and negative reactions). The QC of stains and reagents were maintained by preparing a control smear and staining it with the stains and reagents to be checked. Control strains of *P aeruginosa* (ATCC 27853), *S aureus* (ATCC 25923), *E coli* (ATCC 25922) were used for the standardization and correct interpretation of zone of inhibition of antibiotics during Antibiotic susceptibility testing.

### 2.5 Data interpretation

On the basis of antibiotic susceptibility testing, organism’s resistant to 3 or more than 3 different classes of antibiotics were considered as MDR organisms and interpretation was made as accordingly. Different isolates and antibiotics sensitivity pattern were expressed in percentage by the help of Excel 2016.

## 3. RESULT

In this research total 140 environmental samples were taken, among them, 100 swabs from the various medical-devices’ surfaces were taken and 25 air samples were taken from different locations and 15 swabs from housekeeping surfaces were taken. Of the total environmental samples taken from 17 different sampling sites were considered (appendix H). All samples were found to be growth positive n=140(100%).

### 3.1 Microbial load in air of different location

A total of 25 air samples collected from the different operation theaters and wards by plate exposure method, revealed the growth positivity in all the samples shown in Table 1. It was found that surgical ward had highest number of known air flora, followed by orthopedic ward and medical ward with equal number, then gynae ward 15 emergency room and SICU 14 with equal number, POW 12 and Major OT 12 and least air flora at Minor OT 5.

**Table 1:** Microbial load of isolates in the sample of air circulated.

| Microorganisms | *S. aureus* | *CoNS* | *Micrococci spp* | *Streptococcus spp* | *Bacillus spp* | *Pseudomonas spp* | *Enterobacteriaceae member* | *Fungi* | Total |
| --- | --- | --- | --- | --- | --- | --- | --- | --- | --- |
| Major OT (n=5) | 2 | 3 | 2 | - | 1 | 1 | - | 3 | 12 |
| Minor OT (n=5) | - | 2 | - | - | 2 | - | - | 1 | 5 |
| SICU (n=5) | 3 | 3 | 2 | 1 | 2 | - | 1 | 2 | 14 |
| POW (n=5) | - | 4 | 3 | - | 2 | 1 | - | 2 | 12 |
| Medical ward (n=1) | 2 | 3 | 3 | 1 | 3 | 1 | 2 | 3 | 18 |
| Surgical ward (n=1) | 3 | 4 | 2 | 3 | 4 | 2 | 3 | 1 | 22 |
| Gynae ward (n=1) | - | 5 | - | 2 | 3 | - | 2 | 3 | 15 |
| Orthopedic ward (n=1) | 3 | 2 | 3 | - | 3 | 2 | 2 | 1 | 16 |
| Emergency<br>(n=1) | - | 4 | 1 | 3 | 2 | 1 | 2 | 1 | 11 |

*n=Number of times the sampling was taken.*

### 3.2 Microbial isolates in the surface swab of housekeeping surfaces

A total of 15 swab samples collected from the different operation theaters and wards revealed the growth positivity in all the samples (100%) shown in Table 2. It was found that *S. aureus*, CoNS, *Micrococcus* spp, *Streptococcus* spp, *Bacillus* spp, *Pseudomonas* spp and enterobactereace member were present in all wash basins sampled. Whereas, CoNS, *Micrococcus* spp, *Bacillus* spp were found in all floor swabs.

**Table 2:**
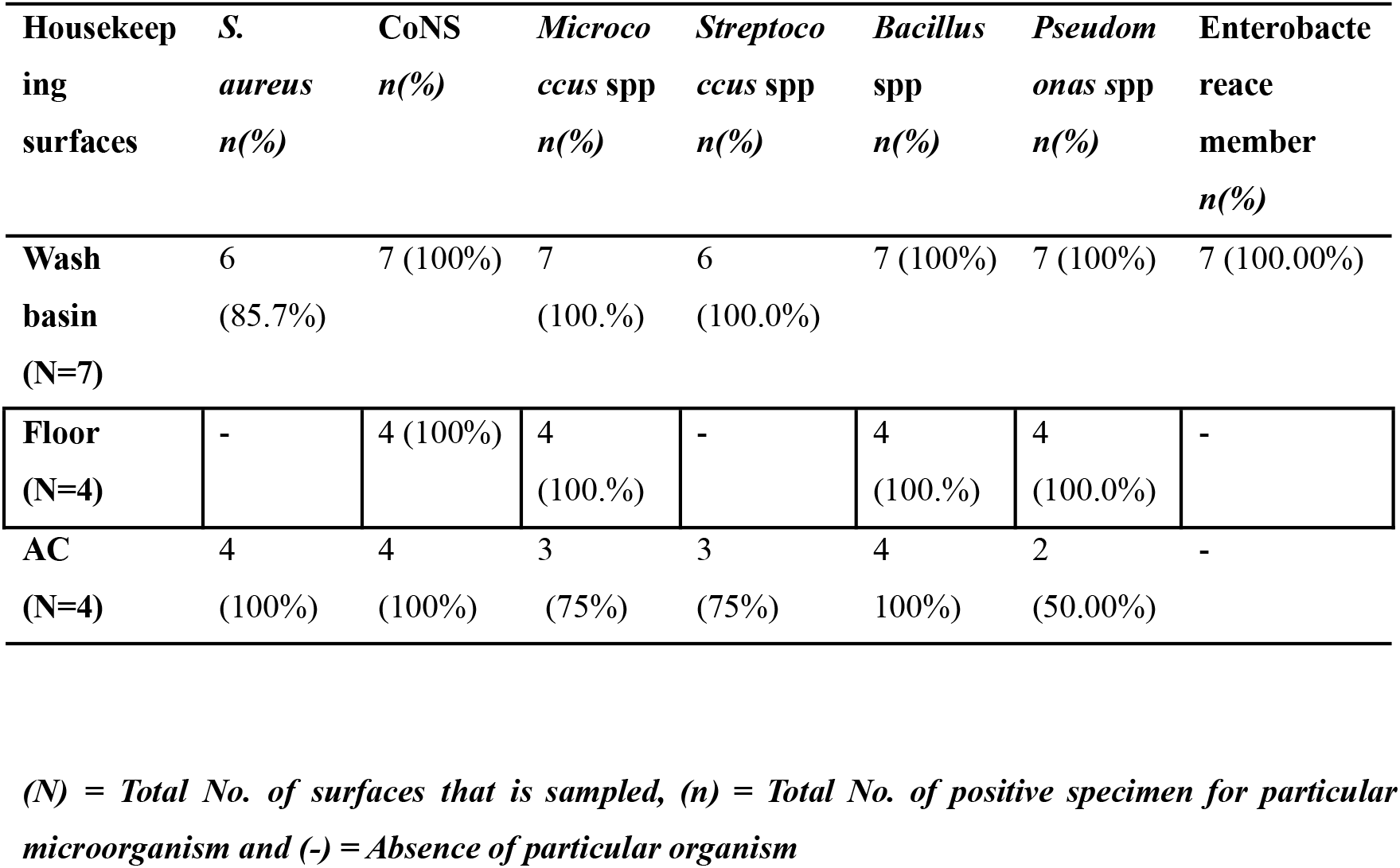
Microbial isolates in the surface swab of housekeeping surfaces.

### 3.3 Microbial isolates in the surface swab of Medical-device surfaces

In the 100 swabs taken from the medical devices from various operation theaters, wards and OPDs. 100% samples showed the growth positivity even the autoclaved instruments (scissors, speculum, forceps and dental instruments) shown in Table 3. The autoclaved instruments showed the growth of CoNS, *Micrococcus* spp and *Bacillus* spp,

**Table 3:**
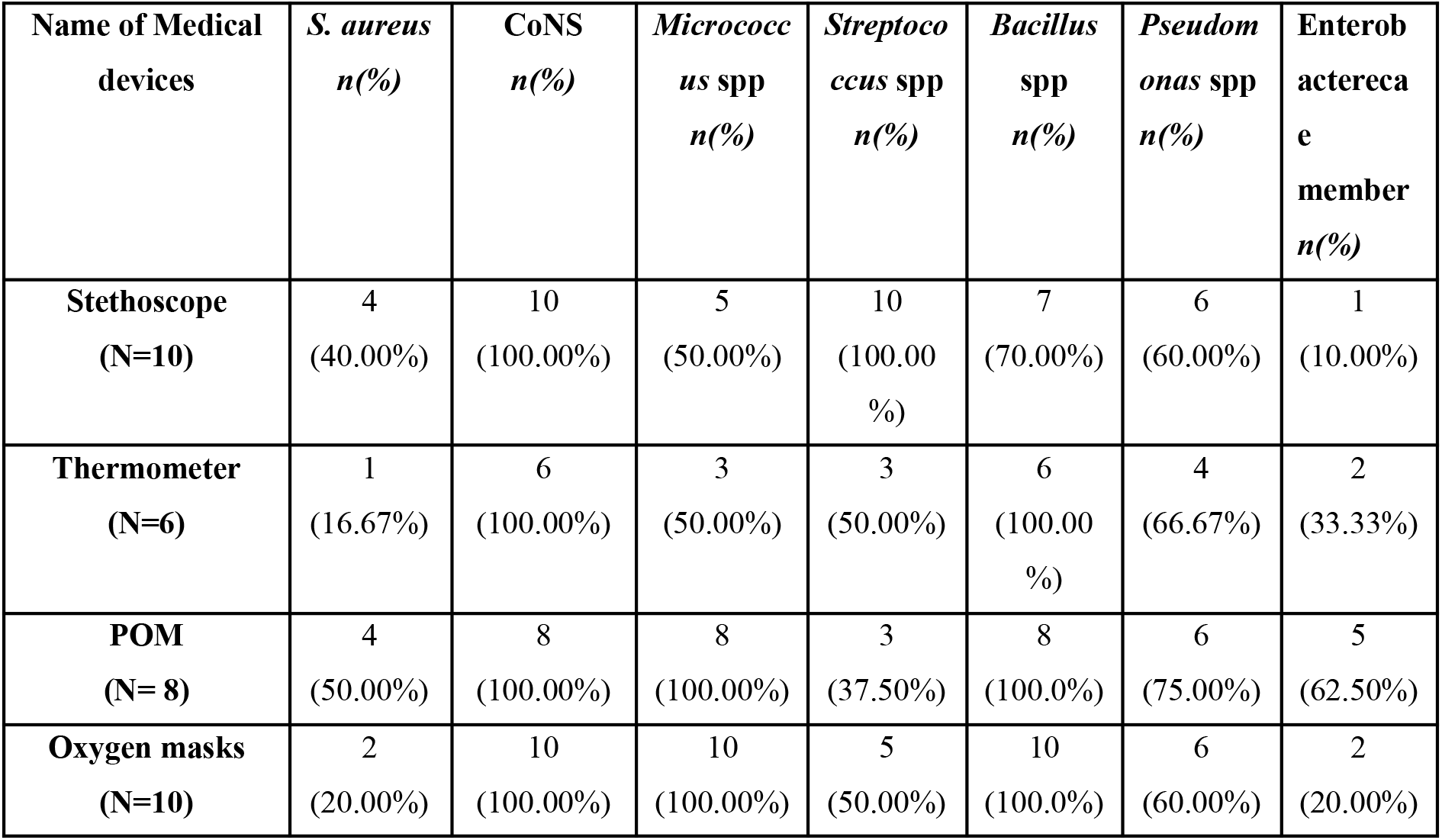
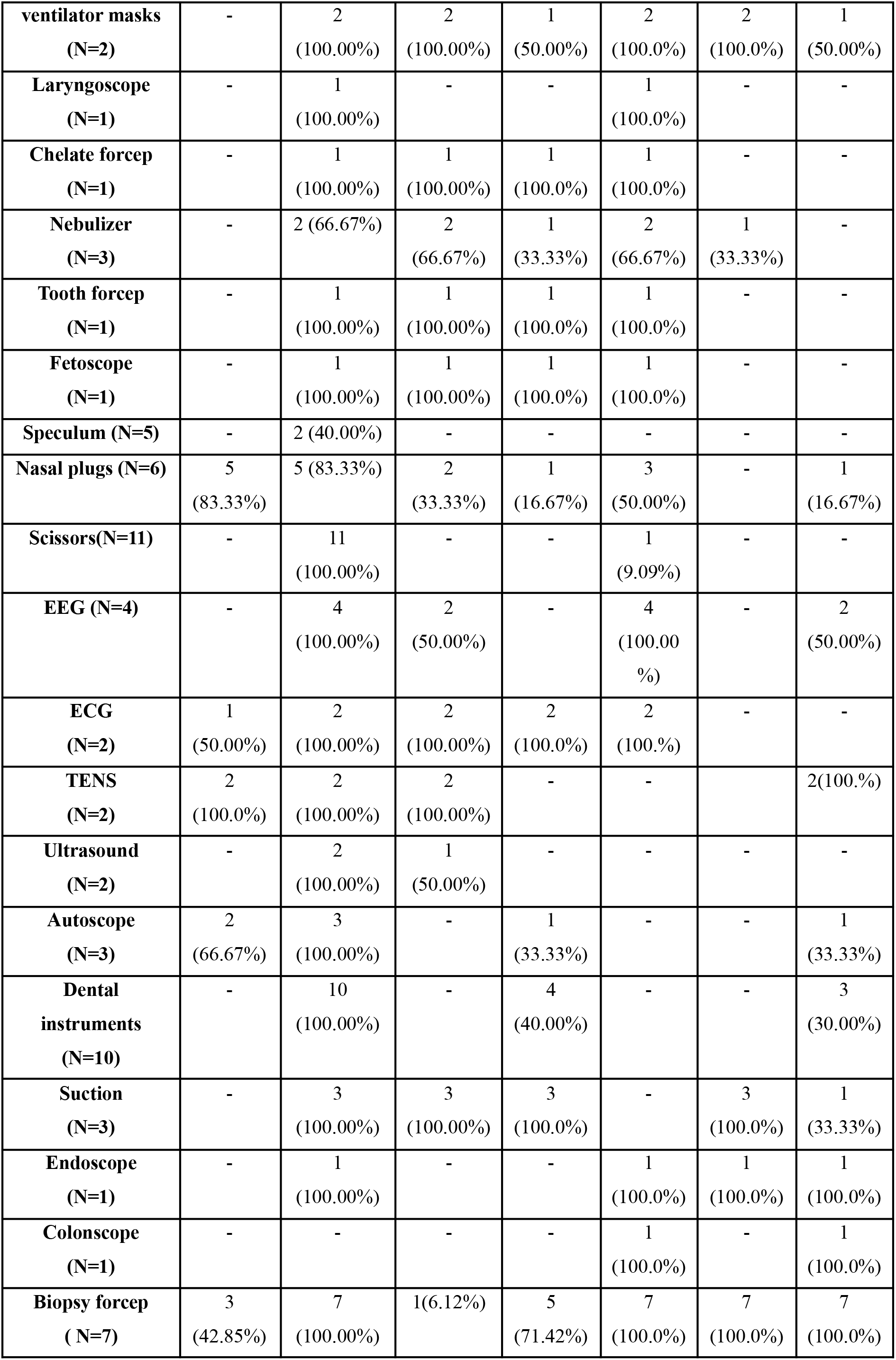
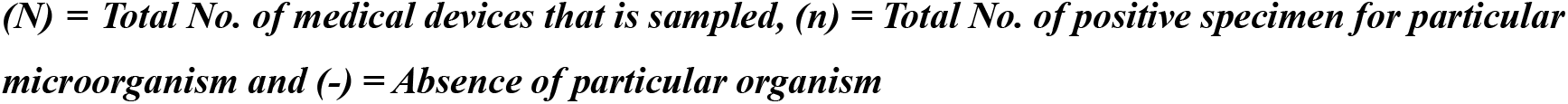
Microbial isolates in the surface swab of medical-device surfaces.

The devices like endoscope, colonoscope and biopsy forceps were disinfected by cidex showed the high number of members of Enterobacteriaceae and *Pseudomonas* spp unlike in other devices which are predominantly contaminated with CoNS, *S. aureus*, *Micrococcus* spp, *Streptococcus* spp, *Bacillus* spp,

### 3.4 Antibiotic Resistivity pattern of the environmental isolates

Altogether 277 Gram positive and Gram negative isolates were subjected to AST.

#### 3.4.1 Antibiotic sensitivity shown by Gram positive isolates

Most of the *Staphylococcus aureus* isolates were sentitive against Cefoxitin 24 (88.89%) and most resistant towards Ofloxacin 24 (88.89%).In case of CONS, most of isolates were sensitive to Gentamicin 34 (73.91%) whereas resistant towards Amoxicilin 25 (54.35%). *Micrococcus* spp was most sensitive towards Ceftriaxone 40 (93.02%) and most resistant against Co-Trimoxazole 18 (41.86%). Similarly, *Streptococcus* spp was most sensitive towards Cefoxitin 30 (96.77%) and resistant to Amoxcilin 4 (62.79%).

#### 3.4.2 Antibiotic sensitivity pattern of Gram negative isolates

Most of the *Pseudomonas* spp isolates were sentitive against Co-Trimoxazole 16 (72.73%) antibiotic and least sensitive towards Ciprofloxacin 11 (50.50%). In case of *E coli*, most of isolates were sensitive to Co-Trimoxazole 28(96.55%) and were resistant towards gentamicin 20 (68.97%). *Acinetobacter* spp was most sensitive towards Co-Trimoxazole 18 (94.74%) and resistance with Nitrofurantoin 17 (89.05%). *Klebsiella* spp was most sensitive towards Ceftriaxone 16 (88.89%) and resistant to gentamicin 5 (76.47%) shown in Table 5. Likewise, the *Proteus* spp was sensitive to Ciprofloxacin 3 (100.00%) and resistant to Ceftriaxone 3 (100.00%) used.

**Table 5:**
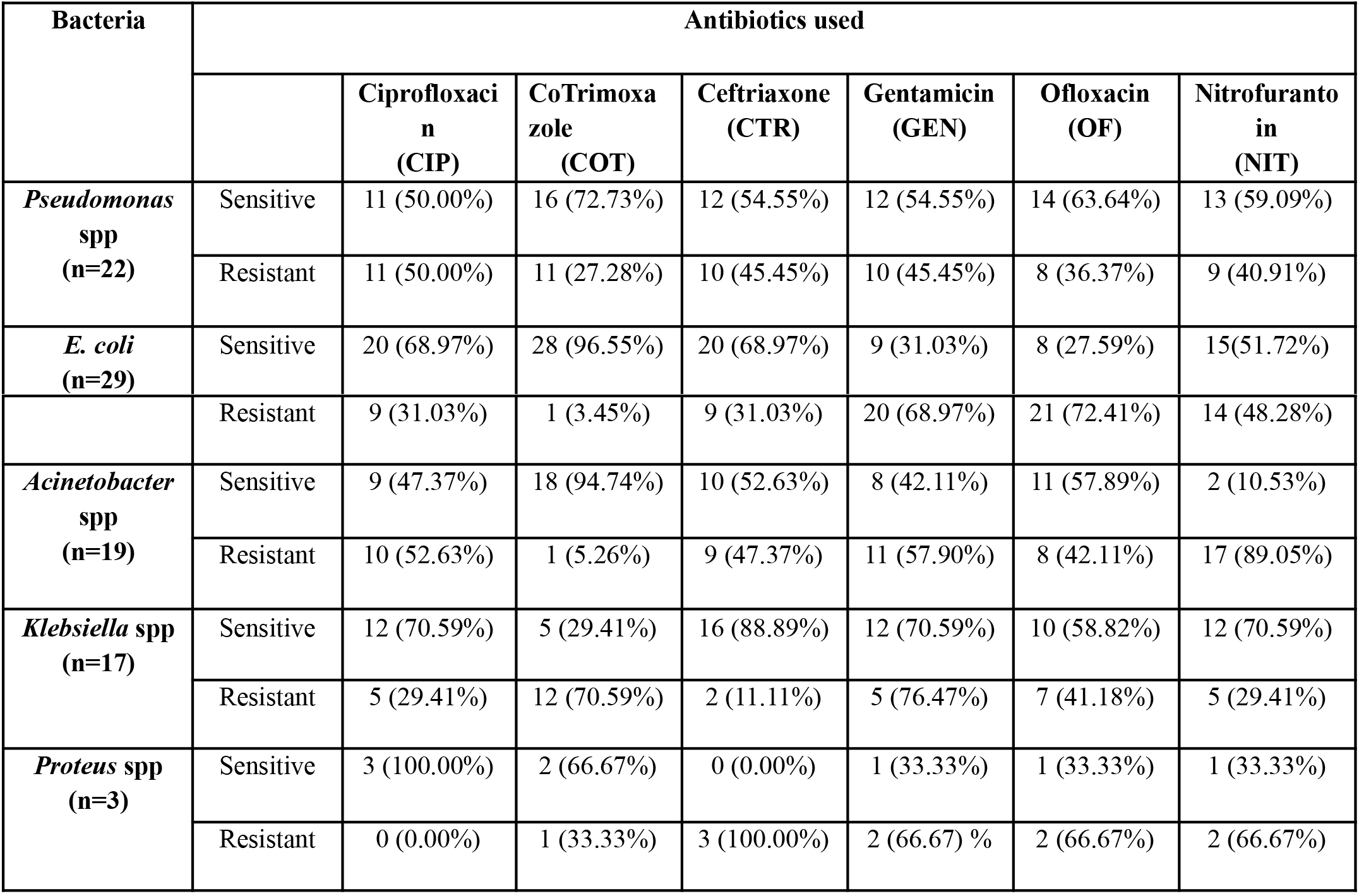
Antibiotic sensitivity shown by Gram negative isolates.

## 4. DISCUSSION

Indoor hospital surfaces are contaminated by frequent contact with patients and health care workers. Despite the various hospital infection control measures, number of data shows that environmental contamination plays an important role in the spread of possible pathogenic bacteria. Environmental contamination may play a key role in the occurring and transmission of NIs, when health care workers contaminate their hands or gloves by touching contaminated surfaces or when patients come into direct contact with contaminated surfaces. So, its assessment is very necessary. Most of these pathogens may remain viable in the environmental aspects of hospitals like air, dust, clothes and in inanimate surfaces and equipment, which can serve as important source of pathogens particularly to the immunocompromised patients (Neely et al 2000 and 2001; Hota et al 2004; Kramer et al 2006).

Total 140 samples were considered, that encompasses the medical devices of hospital (n=100), housekeeping surfaces like floor, wash basin and Air Conditioner (n=15) and air (n=25). Furthermore, antibiotics susceptibility pattern of the isolates was also done.

From the total environmental samples (140) In most of the swab taken, Coagulase Negative Staphylococci was dominant, followed by *Staphylococcus aureus, Streptococcus* spp. *Micrococcus* spp., *E coli, Pseudomonas* spp., *Bacillus* spp., *Acinetobacter* spp., *Klebsiella* spp., Fungi, and least *Proteus* spp. Similar result was obtained in the study done by Pokharel et al (1993), Banjara et al (2002), Panta et al (2006), Sharma et al (2006), Karki et al (2010) and Panta et al (2011) indicating the dominance of gram positive cocci, then gram positive bacilli and finally gram negative bacilli. This was similar to the findings of a study carried by Sapkota et al (2016) in which *S aureus*, *Micrococcus* spp, coagulase negative *Staphylococcus* (CONS), *Bacillus* spp, *Pseudomonas* spp, yeast, and *Acinetobacter* spp were the most commonly detected organisms.

Whereas, gram negative MDR organisms in hospital environment shows that *Klebsiella spp* 6 (35.29%) had highest MDR prevalence followed by *Acinetobacter* spp 6 (31.57%). E. *coli* 8 (27.58%), *Pseudomonas* spp 4 (18.18%) and lastly *Proteus* spp had no MDR at all. Similarly, in the study conducted by Panta (2011) it was found that 109 out of 154 *Acinetobacter* spp (70.8%) was found to be MDR, 13 out of 15 *E. coli* (86.7%) isolates were MDR and among the isolated *Klebsiella* spp All isolates (100.0%) found to be MDR which is not in consistent with results obtained in this research.

There are no any criteria to say any surface as “Clean” by using the aerobic colony counts (Sehulster et al 2003; Dancer 2004) and additionally there is no defined level of microbial load that correlates with hygiene.

## 5. CONCLUSION AND RECOMMENDATIONS

### 5.1 Conclusion

This study was focused to evaluate microbiological quality of air and medical device surfaces of hospital. Total 140 samples were analyzed, that encompasses the medical devices of hospital (n=100), housekeeping surfaces like floor, wash basin and Air Conditioner (n=15) and air (n=25). From this study of microbiological analysis of various isolates from environmental samples, it is found that the CoNS were major contaminant of the hospital. followed by *S. aureus, Streptococcus* spp., *Micrococcus* spp. *E. coli, Pseudomonas* spp., *Bacillus* spp., *Acinetobacter* spp., *Klebsiella* spp., Fungi, and lastly *Proteus* spp. Multi drug resistant bacteria was found in all isolates except *Bacillus* spp. and *Proteus* spp. Hospital environment and medical devices were found to be heavily contaminated with microorganisms even containing thick layer of dust covering it. Hence, proper sanitation, regular monitoring and disinfection are needed to prevent NIs.

### 5.2 Recommendations

1. Based on the study following recommendations have been made: -
2. Disinfection of surfaces of medical devices after every single use must be strictly maintained
3. Fumigation of air of OT is required regularly.
4. Formation of Infection Control Committee (ICC) as per WHO guideline 2004 is required to monitor the possible outbreak of NIs.
5. Wearing masks and hand wash practice is highly recommended.Isolation room required for previous patients infected or colonized with multidrug-resistant (MDR) organisms

## Supporting information

https://docs.google.com/document/d/1-qFOmRF5X3O-8r3NQrwtwiVhGaHmoZxnGPYrEPuQEIQ/edit

## ABBREVIATIONS

AC: Air Conditioner
AST: Antibiotic Sensitivity Test
BA: Blood Agar
CDC: Centers for Disease Control
CLSI: Clinical Laboratory Standard Institute
Cm: Centimeter
CoNS: Coagulase Negative Staphylococci
CRI: Catheter Related Infections
ECG: Electrocardiogram
EEG: Electroencephalogram
ENT: Ear, Nose and Throat
HCW: Health Care Worker
ICU: Intensive Care Unit
KNFH: Korea Nepal Friendship Hospital
MA: MacConkey Agar
MDR: Multi Drug Resistant
MHA: Muller Hinton Agar
MRSA: Methicillin Resistant *Staphylococcus aureus*
NA: Nutrient Agar
NI: Nosocomial infection
NNISS: National Nosocomial Infection Surveillance system
OT: Operation Theater
OPD: Outdoor patient department
POM: Pulse Oxy Meter
POW: Post-Operative Ward
SDA: Sabourd Dextrose agar
SICU: Surgical Intensive Care Unit
SIM: Sulphide Indole Motility
Sq: Square
TSI: Triple Sugar Iron
UTI: Urinary Tract Infection
WHO: World Health Organization

---

**Figure 2:**
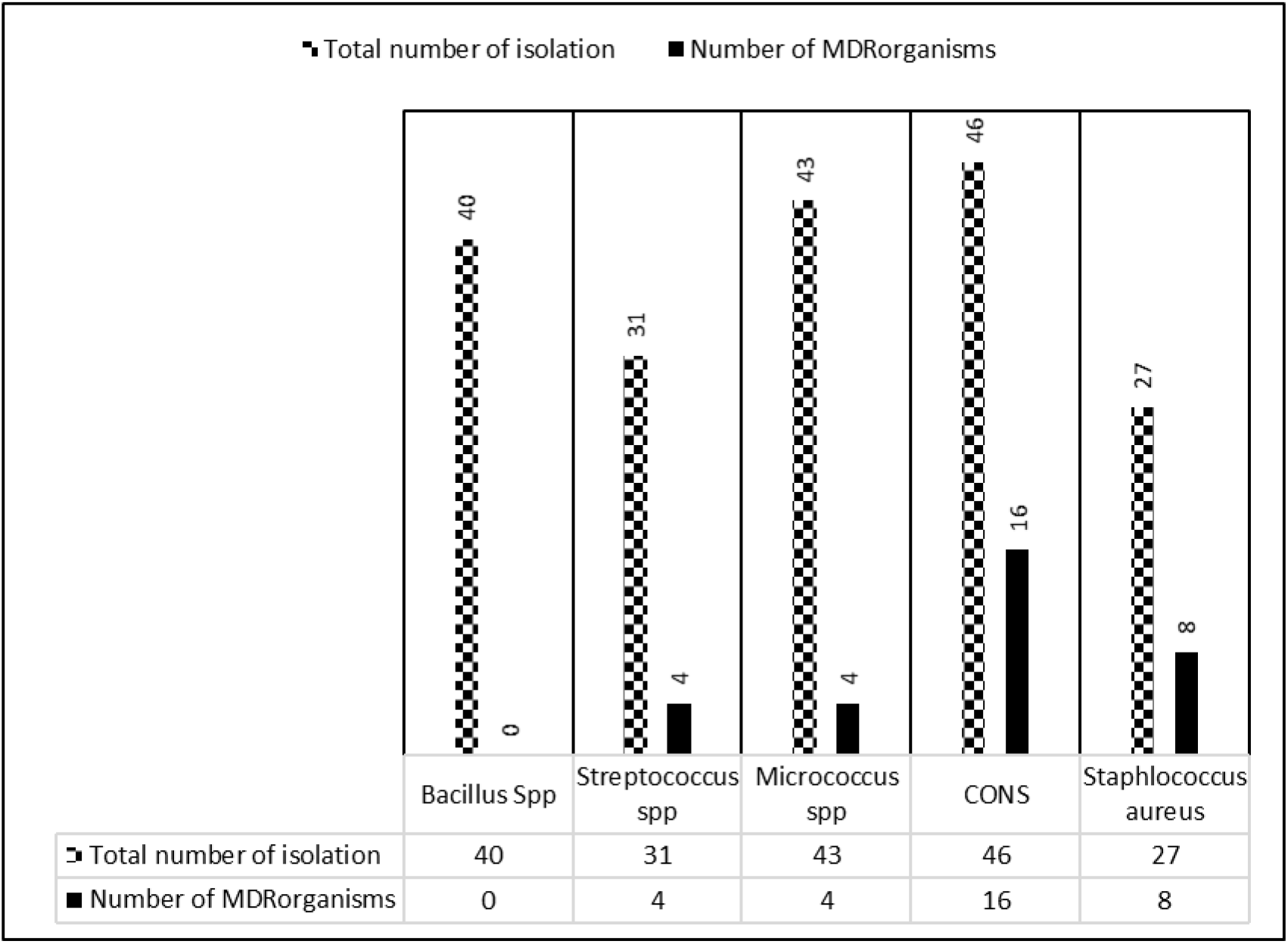
Gram positive MDR organisms. The graph shows that CONS 16 (34.78%) had highest MDR prevalence followed by *Staphylococcus aureus* 8 (29.62%), *Streptococcus* spp 4 (12.90%), *Micrococcus* spp 4 (9.30%) and no MDR was found *Bacillus* spp .

**Figure 3:**
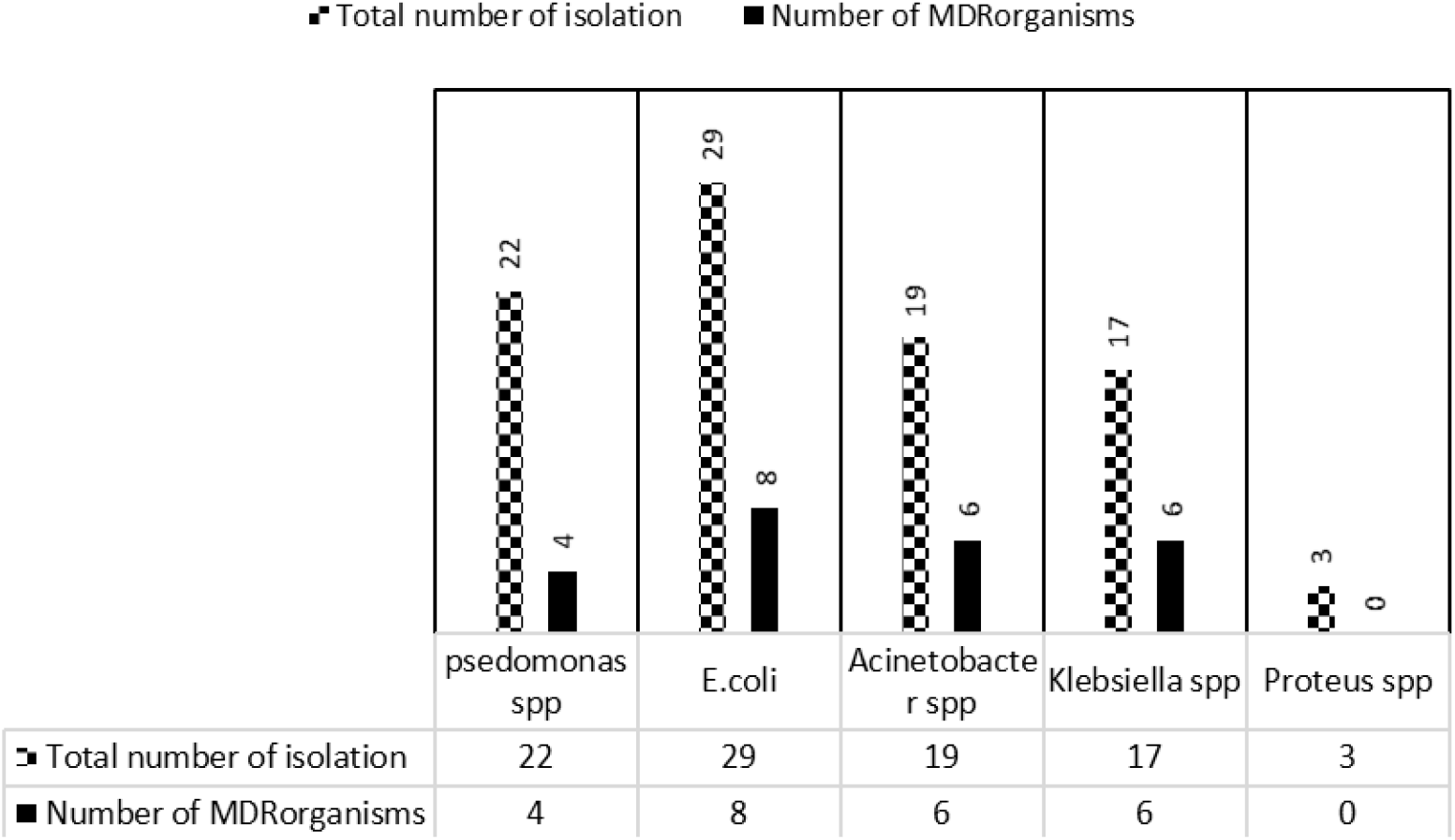
Gram negative MDR organisms. The graph shows that *Klebsiella* spp 6 (35.29%) had highest MDR prevalence followed by *Acinetobacter* spp 6 (31.57%). *E coli* 8 (27.58%) *Pseudomonas* spp 4 (18.18%) and *Proteus* spp had no MDR at all.

