## Supplementary material for "Prevalence Of Multi Drug Resistant Bacteria On Environmental And Medical Device Surfaces Of Korea Nepal Friendship Hospital": https://docs.google.com/document/d/1-qFOmRF5X3O-8r3NQrwtwiVhGaHmoZxnGPYrEPuQEIQ/edit: supplementary.docx

**APPENDICES**

**APPENDIX-A**

**List of materials used in study: -**

**A. Equipments and materials used during the study**

Autoclave	ALP Co. LTD.

Hot air oven	Memmert

Incubator	Memmert

Refrigerator	LG (Korea )

Microscope	Olympus (Japan)

Weighing Machine	Tanita (Japan)

**APPENDIX-B**

**A. Composition and Preparation of Different Culture Media**

**The culture media used were from HiMEDIA company.**

**1.	Nutrient Agar (NA)**

Composition	gram/litre

Peptic digest of animal tissue	5.00

Beef Extract	1.50

Yeast Extracts	1.50

Sodium Chloride	5.00

Agar	15.00

Final pH (at 25°C)	7.4 ± 0.2

28 gm of the medium was suspended in 1000ml of the distilled water and boiled to dissolve completely. Then medium was autoclaved at 121°C (15lbs pressure) for 15min. The sterilized medium was then poured into the sterilized petridishes and then was allowed to cool.

**2.	Blood Agar Base (Infusion Agar)**

Composition	gm/litre

Beef heart, infusion from	500

Tryptose	10.0

Sodium Chloride	5.00

Agar	15.0

Final pH (at 25°C)	7.3 ± 0.2

40gms of the medium was suspended in 1000ml of the distilled water, dissolved by boiling and sterilized by autoclaved at 121°C (15lbs pressure) for 15minutes. After cooling to 50^o^C, 5%v/v sterile defibrinated blood was added aseptically, then mixed with gentle rotation and poured into the sterilized petridishes and was allowed to cool.

**3.	MacConkey Agar (MA)**

Composition	gm/litre

Peptic digest of animal tissue	17.00

Proteose Peptone	3.00

Lactose	10.00

Bile Salt	1.50

Sodium Chloride	5.00

Neutral Red	0.03

Agar	15.00

Final pH (at 25°)	7.1 ± 0.2

51.53gm of the medium was dissolved in 1000ml of the distilled water and then boiled to dissolve completely. The media was autoclaved at 121°C (151bs pressure) for 15min. The sterilized medium was then poured in sterilized petridishes and was allowed to cool.

**4.	Nutrient Broth (NB)**

Composition	gm/litre

Peptic digest of animal tissue	5.00

Sodium Chloride	5.00

Beef Extract	1.50

Yeast Extracts	1.50

Final pH (at 25°)	7.4 ± 0.2

13gm of the medium was dissolved in 1000ml of the distilled water and then boiled to dissolve completely. The medium was then dispensed in test tube in amount of 3ml in each and autoclaved at 121°C (151bs pressure) for 15 minutes. The sterilized medium was then cooled to room temperature.

**5.	Muller Hinton Agar (MHA)**

Composition	gm/litre

Beef infusion form	300.0

Casein Acid Hydrolysate	17.50

Starch	1.50

Agar	17.00

Final pH (at 25°)	7.3 ± 0.2

38gm of the medium was dissolved in 1000ml of the distilled water and then boiled to dissolve completely. The medium was autoclaved at 121°C (15lbs pressure) for 15minutes. The sterilized medium was then poured in sterilized petridishes and was allowed to cool.

**6. Mannitol Salt Agar (MSA)**

Composition	gm/litre

Proteose Peptone	10.00

Beef Extract	1.000

Sodium Chloride	75.00

D-Mannitol	10.00

Phenol Red	0.025

Agar	15.00

Final pH (at 25°)	7.4 ± 0.2

111gm of the medium was dissolved in 1000ml of the distilled water and then boiled to dissolve completely. The medium was autoclaved at 121°C (15lbs pressure) for 15minutes. The sterilized medium was then poured in sterilized petridishes and was allowed to cool.

**7. MacConkey Broth Purple (for MPN method)**

Composition	gm/litre

Peptic digest of animal tissue	20.0

Lactose	10.0

Sodium Chloride	5.00

Sodium Taurocholate	5.00

Bromocresol Purple	0.01

Final pH (at 25°)	7.4 ± 0.2

40gm of the medium was dissolved in 1000ml of the distilled water and then boiled to dissolve completely. The medium was then distributed in test tubes with inverted Durham’s tube and autoclaved at 121°C (151bs pressure) for 15 minutes. The sterilized medium was then cooled to room temperature.

**B.	Composition and preparation of different biochemical media**

**1. Simmon Citrate Agar**

Composition	gm/litre

Magnesium Sulfate	0.20

Monoammonium Dihydrogen Phosphate	1.00

Dipotassium Phosphate	1.00

Sodium Citrate	2.00

SodiumChloride	5.00

Agar	15.0

Bromothymol Blue	0.08

Final pH (at 25°C)	6.8 ± 0.5

24.2 grams of the medium was dissolved in 1000ml of the distilled water and boiled to dissolve completely. 3 ml of the medium was dispensed in each test tube and autoclaved at121°C (15lbs pressure) for 15minutes. The sterilized medium in the test tube was then allowed to set in slopes or slant.

**2. Urea Agar base (Christensen urea agar)**

Composition	gm/litre

Peptic digest of animal tissues	1.000

Dextrose	1.000

Monopotassium Phosphate	0.800

Dipotassium Phosphate	1.200

Sodium Chloride	5.000

Agar	15.00

Phenol Red	0.012

Final pH (at 25C)	6.8 ± 0.2

24 grams of the medium was suspended in 950 ml of distilled water, dissolved by boiling and autoclaved at 121^0^C (15 lbs pressure) for 15 minutes. After cooling to 50^0^C, 50 ml of sterile 40% urea solution was added aseptically, mixed with gentle rotation. Then 5ml of the medium was dispensed in test tube and set at slant position.

**3. Sulphide Indole Motility (SIM) Agar**

Composition	gm/litre

Peptic digest of animal	30.00

Beef Extract	3.00

Peptonized Iron	0.20

Sodium Thiosulfate	0.025

Agar	3.00

Final pH (at 25°C)	7.3 ± 0.2

36.23grams of the medium was dissolved in 1000ml of the distilled water, boiled to dissolve completely and then dispensed in test tubes to a depth of about 3 inches. Then the medium in tubes was autoclaved at121°C (15lbs pressure) for 15 minutes.

**3. MR-VP Medium**

Composition	gm/litre

Buffered peptone	7.00

Dextrose	5.00

Di-potassium phosphate	5.00

Final pH (at 25°C)	6.9 ± 0.2

17 grams of the medium was dissolved in 1000ml of the distilled water, boiled to dissolve completely and then 3ml of the medium was dispensed in each test tubes and then autoclaved at121°C (15lbs pressure) for 15 minutes.

**4.	Triple Sugar Iron (TSI) Agar**

Composition	gm/litre

Peptic digest of animal tissue	10.00

Casein Enzymatic Hydrolysate	10.00

Yeast Extracts	3.00

Beef Extract	3.00

Lactose	10.00

Sucrose	10.00

Dextrose	1.00

Sodium Chloride	5.00

Ferrous Sulphate	0.20

Sodium Thiosulphate	0.30

Agar	12.00

Phenol red	0.024

Final pH (at 25°C)	7.4 ± 0.2

65grams of the medium was dissolved in 1000ml of the distilled water. The medium was then dispensed in test tubes and autoclaved at 121°C (15lbs pressure) for 15minutes. The sterilized medium in the test tube was then allowed to set in slant with a butt of about 1 inch of thickness.

**C. Composition and preparation of different staining reagent**

**1. Gram Stain**

**(a) Crystal Violet Solution**

Crystal Violet	20.0 g

Ammonium Oxalate	9.0 g

Ethanol or Methanol	95.0 ml

Distilled Water (D/W) to make	1 litre

***Preparation:*** 20 grams of Crystal Violet was weighed in a clean piece of paper and transferred to a clean brown bottle. Then 95 ml of ethanol was added and mixed until the dye is completely dissolved. To the mixture, 9 grams of Ammonium Oxalate dissolved in 200ml of D/W was added. Finally the volume was made 1 litre by adding D/W.

**(b) Lugol's Iodine**

Potassium Iodide	20.00g

Iodine	10.00g

Distilled Water	1000.0 ml

***Preparation:*** To 250 ml D/W, 20 grams of Potassium Iodide was dissolved. Then 10 grams of iodine was mixed to it until it was dissolved completely. Finally the volume was made 1 litre by adding D/W.

**(c) Acetone**-**Alcohol Decolorizer**

Acetone	500ml

Ethanol (absolute)	475ml

Distilled Water	25.0 ml

***Preparation:*** 475 ml of ethanol (absolute) was added to 25ml of D/W, mixed and transferred into a clean bottle. Then immediately, 500ml of acetone was added to the bottle and mixed well.

**(d) Safranin (Counter Stain)**

Safranin (2.5% solution in

95% ethyl alcohol)	10.00ml

Distilled Water	100.0 ml

***Preparation:*** 2.5 % of Safranin solution was prepared in 95% ethanol. 10 ml of this solution was then suspended in 100 ml of D/W.

**2. Normal saline**

Sodium Chloride	0.85g

Distilled Water	100ml

***Preparation:*** 0.85 grams of Sodium Chloride was weighed and added to a bottle containing 100ml of D/W and mixed well to dissolve the salt completely. The bottle was well labeled and stored at room temperature.

**3. Biochemical Test Reagents**

**a. For catalase test**

**Catalase reagent (3%H2O2)**

Hydrogen Peroxide	1ml

Distilled Water	9ml

***Preparation:*** To 9ml of D/W, 1ml of Hydrogen Peroxide was added and mixed well so as to make a 3% solution of Hydrogen Peroxide.

**b. For oxidase test**

**Oxidase reagent (impregnated in Whatman’s No. 1 filter paper)**

Tetramethyl *p*-phenylene diamine dihydrochloride(TPD)	1.00g

Distilled Water	100ml

***Preparation:*** 1 gram of TPD was dissolved in 100 ml of D/W. To this solution, strips of Whatman’s No.1 filter paper were soaked and drained for about 30 seconds. Then these strips were freeze dried and stored in a dark bottle tightly sealed with a screw cap.

**c. For indole test**

**Kovac's Indole Reagent**

*p*-dimethyl aminobenzyldehyde	2.00gm

Isoamyl alcohol	30.0ml

Concentrated Hydrochloric Acid	10.0ml

***Preparation:*** In 30 ml of isoamyl alcohol, 2 grams of *p*-dimethyl aminobenzaldehyde was dissolved and transferred to a clean brown bottle. Then to this solution, 10ml of concentrated Hydrochloric Acid was added and mixed well.

**d. For methyl red test**

**Methyl red solution**

Methyl red	0.05gm

Ethylalcohol(absolute)	28.0ml

Distilled Water	22.

***Preparation:*** 0.05 gram of methyl red was dissolved in 28 ml ethanol and transferred to a clean brown bottle. To this, 22 ml of D/W was added and mixed well.

**e. For Voges Proskauer test**

**Barrit’s reagent**

Solution A

α-Napthol	5.0gm

Ethyl alcohol (absolute)	100ml

***Preparation:*** 5 gram of α napthol was dissolved in 25 ml ethanol and transferred into a clean brown bottle.Then the finel volume was made 100ml by adding ethanol.

Solution B

Potassium hydroxide (KOH)	40.0gm

Distilled Water	100ml

***Preparation:*** 40 gram of KOH was dissolved in 25 ml D/W and transferred into a clean brown bottle. Then the final volume was made 100 ml by adding D/W.

**APPENDIX-C**

**A. Procedure for Gram Staining (Forbes *et al*., 2007)**

Gram staining is a differential staining that differentiates all bacterial species into two large groups: *gram positive* and *gram negative*. The following steps were involved in gram staining:

1. A thin film of the material to be examined was prepared and dried.
2. The material on the slide was heat fixed and allowed to cool before staining.
3. The slide was flooded with Crystal Violet stain and allowed to remain without
4. drying for 10-30 seconds.
5. The slide was rinsed with tap water, shaking off excess.
6. The slide was flooded with iodine solution and allowed to remain on the
7. surface without drying for twice as long as the crystal violet was in contact
8. with the slide surface.
9. The slide was rinsed with tap water, shaking off excess.
10. The slide was flooded with alcohol acetone decolorizer for 10 seconds and
11. rinsed immediately with tap water until no further colour flows from the slide
12. with the decolorizer. Thicker smear requires more aggressive decolorization.
13. The slide was flooded with counter stain (safranin) for 30 seconds and washed
14. off with tap water.
15. The slide was blotted between two clean sheets of bibulous paper and

examined microscopically under oil immersion at 100X

**APPENDIX-D**

**Methods of biochemical test used for the identification of pathogens**

**a. Catalase test**

The enzyme Catalase is present in most cytochrome containing aerobic and facultative anaerobic bacteria; the main exception is Streptococcus species (catalase negative). Usually organisms which lack the cytochrome system also lack the Catalase enzyme and therefore are unable to break down hydrogen peroxide. Catalase is a heme protein. The prosthetic group is made up of four atoms of trivalent iron (ferric) per molecule, which retains its oxidized state during enzyme activity. Hydrogen peroxide is formed as an oxidative end product of the aerobic breakdown of sugars. Reduced flavoprotein reacts directly with gaseous oxygen by way of electron reduction to form hydrogen peroxide and not by direct action between hydrogen and molecular oxygen.

With the help of a sterile glass rod, a small amount of culture from the Nutrient Agar was transferred to a clean glass slide and a drop of 3% Hydrogen peroxide solution was dropped on the surface of the slide. The positive test is indicated by the formation of active bubbling of the oxygen gas. The lack of Catalase was evident by lack of or weak bubble production.

**b. Oxidase test**

The Oxidase test is based on the bacterial production of an Oxidase enzyme. The oxides reaction is due to the presence of a cytochrome oxidase system which activates the oxidation of reduced cytochrome by molecular oxygen, which in turn acts as an electron acceptor in the terminal stage of the electron transport system.The cytochrome system is usually present only in aerobic organisms which make them capable of utilizing oxygen as a final hydrogen acceptor to reduce molecular oxygen to hydrogen peroxide, the last link in the chain of aerobic respiration.

A piece of filter paper soaked in Oxidase reagent and dried was moistened with distilled water and a colony from the fresh culture was picked up with a sterile glass rod and smeared on the paper. The positive test is indicated by the appearance of blue-purple color within 10 seconds. The Oxidase reagent (Tetra methyl *p*- phenylene diamine dihydrochloride) is a dye that is primary aromatic amines and diamine derivatives of benzene. Cytochrome oxidase in the presence of atmospheric oxygen oxidizes the Oxidase reagent to form a colored compound called indophenol. The Oxidase test is based on the bacterial production of an Oxidase enzyme.

**c. Indole production test**

Tryptophan is an amino acid that can be oxidized by certain bacteria to form three major indolic metabolites: indole, skatol (methyl indole), and indole acetic acid. Intracellular enzymes involved in this oxidation process are collectively called as Tryptophanase. The enzyme tryptophanase catalyses the deamination reaction attacking the tryptophan molecule in its side chains and leaving the aromatic ring intact in the form of indole.

For this test, organism was stabbed in SIM ( Sulfide Indole Motility ) medium from the nutrient broth and incubated at 37^o^C for24 hours . After incubation, 2-3 drops of Kovac’s reagent (*p-* dimethyl aminobenzaldehyde in acid ethanol) was added and resulting color was noted .Indole, if present combines with the aldehyde present in Kovac’s reagent to give a red color in the alcohol layer . The color reaction is based on the presence of pyrrole structure present in the indole.

**d. Methyl red test**

This test is used to determine the ability of an organisms to produce and maintain the stable acid end product from glucose fermentation, and to overcome the buffering capacity of the system.The methyl red test uses a pH indicator in the form of methyl red, to determine the hydrogen ion concentration (pH) arising out of fermentation of glucose by an organism. The hydrogen ion concentration depends on gas ratio (CO2 and H2), which in turn is an index to the different pathways of glucose metabolism exhibited by various organism. The different fermentation patterns are due to variation in enzymes concerned with pyruvic acid metabolism present in the organism. Methyl red positive organisms produce stable acids, maintaining a high concentration of hydrogen ions until a sudden concentration is reached. The validity of methyl red test depends upon a sufficient incubation period in order to permit the differences in glucose metabolism to occur. The organisms to be tested should be incubated at least 35^o^C -37^o^C. Methyl red is an indicator which is already acidic and well denotes changes in degree of acidity by color reactions over a pH range of 4.4-6.0.A pure colony of the test organism was inoculated into 2 ml of MRVP medium and was incubated at 37^o^C for 24 hours. After incubation, about 5 drops of methyl red reagent was added and mixed well. The positive test was indicated by the development of bright red color, indicating acidity and negative with yellow color.

**e. Voges Proskauer test**

The principle of this test is to determine the ability of some organisms to produce a neutral end product, acetyl methyl carbinol (acetoin), from glucose fermentation.Glucose is metabolized to pyruvic acid which is a key intermediate in glycolysis. From pyruvic acid, there are many pathways that a bacterium may follow. The production of acetoin is one pathway for glucose degradation occurring in bacteria. The VP test for acetoin is used primarily to separate *Escherichia coli* from *Klebsiella* and *Enterobacter* spp.A pure colony of the test organism was inoculated into 2 ml of MRVP medium and was incubated at 37^o^C for 24 hours. After incubation, about 5 drops of Barrit’s reagent was added and mixed well and kept for 15 minutes, positive test shows development of pink red color.

**f. Citrate utilisation test**

This test is done to determine if an organism is capable of utilizing citrate as a sole source of carbon for metabolism with resulting alkalinity .The organism whose ability was to be tested were inoculated in the slant of Simmon’s Citrate agar media and incubated at 37^o^C for 24 hours. Result was interpreted as positive if there was a growth or change in color of slant from green to intense blue and negative if there is no growth and no change in color.

**g. Triple sugar Iron (TSI) Agar:**

Triple sugar Iron (TSI) agar is a medium used in the identification of Gram negative enteric rods. The medium measures the ability of a bacteria to utilize sugars: glucose, sucrose and lactose, the concentration of which are in 0.1%, 1.0% and 1.0% respectively. A pH indicator (Phenol Red) included in the medium can detect acid production from fermentation of test carbohydrates. The test organism was streaked and stabbed on the surface of TSI and incubated at 37^o^C. for 24 hours. The results are interpreted as:

a. Yellow (Acid) / Yellow (Acid), Gas, H2S Glucose, Lactose/ Sucrose fermenter, H2S producer.

b. Red (Alkali) / Yellow (Acid), No Gas, No H2S Glucose fermenter, Lactose/Sucrose nonfermenter, Anaerogenic, H2S nonproducer.

c. Red (Alkali) /No Change	Glucose, Lactose and Sucrose nonfermenter,

d. Yellow (Acid)/ No Change	Glucose oxidizer.

e. No Change / No Change	Nonfermenter.

**h. Motility test:**

Bacteria are motile by means of flagella. Flagella occur primarily among the bacilli; however, a few cocci forms are motile. Motile bacteria may contain single flagella. The motility media used for motility are semi solid, making motility interpretation macroscopic. Motile organisms migrate from the stab- line and diffuse into the medium causing turbidity. They may exhibit fussy streaks of growth. Whereas nonmotile bacteria show the growth along the stab-line, and the surrounding media remains colorless and clear.

**i. Coagulase test**

The Coagulase test is used to especially differentiate species within the genus Staphylococcus. *S. aureus* (usually positive) is differentiated from Staphylococcus saprophyticus and *S. epidermidis* (usually negative). A positive coagulase test is usually the final diagnostic criterion for the identification of Staphylococcus species.

The exact mechanism and chemical structure of coagulase is unknown. It is known, however, that this enzyme plays a thromboplastin like role in clotting by converting fibrinogen to fibrin. In vitro, coagulse increases the rate of plasma clotting, the end result of which is the formation of fibrin clot.

Two types coagulase are produced by most of the *S. aureus*:

A) Free coagulase, which converts fibrinogen to fibrin by activating a coagulase reacting factor present in plasma. It can be detected by the appearance of the fibrin clot in the tube coagulase test.
B) Bound coagulase, also known as clumping factor, converts fibrinogen directly to fibrin without requiring a coagulase reacting factor. It can be detected by the clumping of bacterial cells in the rapid slide coagulase test.

**Slide coagulase test**

For slide coagulase test, a drop of physiological saline as placed on three places of a slide, and then a colony of the test organisms was emulsified in one to make thick suspensions. A drop of plasma was added to the suspensions and mixed gently. Then a clumping was observed within 10 seconds for the positive coagulase test. No plasma was added in second suspension. This is used for the differentiation of any granular appearance of the organism from true coagulase clumping. The third drop of saline aws used for a known strain of coagulase positive staphylococci.

**Tube coagulase test**

For the organism showing positive test on slide coagulase test, tube coagulase test is performed. In the tube coagulase test, 0.5 ml of the diluted plasma (1:10 in physiological saline) was pipetted into 18-24 hours broth culture tube inoculated with test organisms. After mixing gently, the tube was incubated at 37^o^C for 2-6 hours. The clotting is observed by gently tilting the tube for positive coagulase test.

**j. Urea hydrolysis test**

This test demonstrates the urease activity present in certain bacteria which decomposes urea, releasing ammonia. The test organism was inoculated in a medium containing urea and the indicator Phenol red. The inoculated medium was incubated at 37^o^C overnight. Positive organism shows pink red due to the breakdown of urea to ammonia. With the release of ammonia, the medium becomes alkaline as shown by a change in color of the indicator to red pink.

**APPENDIX-E**

**Turbidity standard equivalent to McFarland 0.5**

1% v/v solution of sulphuric acid was prepared by adding 1 ml of concentrated sulphuric acid to 99ml of water. Then to 99.4 ml of this solution, 0.6 ml of 1% w/v solution of barium chloride prepared by dissolving 0.5 gram dihydrate barium chloride (BaCl2.2H2O) in 50 ml of D/W, was added and mixed well. Then the standard was transferred into screw capped tubes of the same size and volume as those used for preparing the test and control inocula. The tubes were then sealed tightly to prevent loss by evaporation and stored protected from light at room temperature. The turbidity standard was then vigorously agitated before use. This standard when stored in well sealed container in the dark at room temperature (20-28°C), may be kept for up to 6 months.

**APPENDIX-F**

**Zone size interpretation chart of antibiotics used in present study**

| **Antibiotics used** | **Symbol** | **Disc potency (mcg)** | **Diameter of zone of inhibition in mm** | |
| --- | --- | --- | --- | --- |
|  |  |  | **Resistant** | **Sensitive** |
| Amoxycillin | AMX | 10 | 13-17 | 20 |
| Azithromycin | AZM | 15 | 13-17 | 18 |
| Ceftriaxone | CTR | 30 | 13-20 | 21 |
| Cotrimoxazole (Trimethoprim+ Sulfamethoxazole) | COT | 1.25+ 23.75 | 10-15 | 16 |
| Gentamicin | GEN | 10 | 12-14 | 15 |
| Cefoxitin | CX | 5 | 9-13 | 14 |
| Nitrofurantoin | NF | 100 | 14-16 | 17 |
| Ofloxacin | OF | 5 | 19-21 | 22 |

(Source: Product Information Guide, HiMedia Laboratories Pvt. Limited, Mumbai, India)

**APPENDIX-G**

**Table 16: - Grouping of antibiotics**

| **Group number** | **Antibiotics Group** | **Antibiotic** |
| --- | --- | --- |
| Group 1 | Aminopenicillins | Ampicillin |
|  |  | Amoxicillin |
| Group 2 | Sulphonomides | co-trimoxazole |
| Group 3 | Quinolones | ciprofloxacin |
| Group 4 | Aminoglycosides | Gentamycin |
|  |  | Amikacin |
| Group 5 | Cephalosporins | Cefotaxime |
|  |  | ceftriaxone |
| Group 6 | Miscellaneous drug | nitrofurantoin |

**APPENDIX-H**

**Collection sites in hospital**

| **Location** | **Collection sites** | **Total number of sites** |
| --- | --- | --- |
| OT(Minor and Major) | Air | 10 |
|  | Wash basin | 2 |
|  | Floor | 4 |
|  | AC | 4 |
|  | Oxygen masks | 3 |
|  | Ventilator masks | 2 |
|  | Stethoscope | 1 |
|  | Laryngoscope | 1 |
| SICU and POW | Air | 10 |
|  | Oxygen masks | 7 |
|  | POM | 4 |
|  | Stethoscope | 2 |
|  | Thermometer | 1 |
| Surgical ward | Air | 1 |
|  | Stethoscope | 1 |
|  | Thermometer | 1 |
|  | POM | 1 |
|  | Chelate forceps | 1 |
|  | scissors | 1 |
| Medical ward | Stethoscope | 1 |
|  | Air | 1 |
|  | Thermometer | 1 |
|  | POM | 1 |
|  | Wash basin | 1 |
|  | nebulizer | 1 |
| Gynae ward and OPD | Stethoscope | 1 |
|  | Air | 1 |
|  | Tooth forceps | 1 |
|  | Fetoscope | 1 |
|  | Speculum | 5 |
| Orthopedic ward | Stethoscope | 1 |
|  | Air | 1 |
|  | Thermometer | 1 |
|  | POM | 1 |
|  | Wash basin | 1 |
| Emergency ward | Stethoscope | 1 |
|  | Air | 1 |
|  | Thermometer | 1 |
|  | POM | 1 |
|  | Nebulizer | 2 |
|  | Nasal plugs | 6 |
|  | scissors | 10 |
| EEG and ECG room | EEG | 2 |
|  | EGG | 4 |
|  | EEG wash basin | 2 |
| Physiotherapy room | Ultrasound | 2 |
|  | TENS | 2 |
| ENT and Dental Unit | Stethoscope | 1 |
|  | Autoscope | 3 |
|  | Dental instruments | 10 |
|  | Suction | 3 |
| Endoscopy room | Endoscope | 1 |
|  | colonoscope | 1 |
|  | Biopsy forceps | 7 |
| Pediatric ward | Stethoscope | 1 |
|  | Thermometer | 1 |
|  | Wash basin | 1 |
| Total | | 140 |
